# Multitrait diversification in marine diatoms in constant and warmed environments

**DOI:** 10.1101/2022.04.08.487611

**Authors:** Jana Hinners, Phoebe A. Argyle, Nathan G. Walworth, Martina A. Doblin, Naomi M. Levine, Sinéad Collins

## Abstract

Phytoplankton are photosynthetic marine microbes that affect food webs, nutrient cycles, and climate regulation. Their roles are determined by a correlated set of phytoplankton functional traits including cell size, chlorophyll content, and cellular composition. Here, we explore how interrelated trait values and correlations evolve. Because both chance events and natural selection contribute to phytoplankton trait evolution, we used population bottlenecks to diversify six genotypes of Thalassiosirid diatoms. We then evolved them in two environments where natural selection could act on this diversity. Interspecific variation and within-species evolution were visualized for nine traits and their correlations using reduced axes (a trait-scape). Shifts in both trait values and correlations, resulting in movement of evolving populations on the trait-scape, occurred in both environments, and were more frequent under environmental change. Which trait correlations evolved was strain-specific, but greater departures from ancestral trait correlations were associated with lower population growth rates. There was no single master trait that could be used to understand multitrait evolution. Instead, repeatable multitrait evolution occurred along a major axis of variation defined by several diatom functional traits and trait relationships. Because trait-scapes capture changes in trait correlations and values together, they offer an insightful way to study multitrait variation.

## Introduction

Phytoplankton functional traits such as cell size, elemental composition, and population growth rates are crucial to the roles that phytoplankton play in ocean ecology and biogeochemical cycling. Functional traits are highly diverse across photosynthetic marine microbes (phytoplankton), allowing them to thrive in all aquatic environments. This diversity evolves both by natural selection and chance events (1). For pelagic marine phytoplankton, this tension between the actions of natural selection and chance is often framed as the roles of local adaptation vs. migration in determining phytoplankton distributions and trait diversity (2–4). However, our understanding of how interrelated functional traits can diversify focuses almost exclusively on adaptation to changing environments, and rarely incorporates variation fixed through chance events. Modeling, predicting, and explaining functional trait evolution requires defining the patterns of viable variation in functional traits generated during chance events, such as founder effects that can occur during migration or at the start of diatom bloom events, since this variation can be what natural selection acts on. Understanding how interrelated functional traits evolve both in the absence of environmental change, and because of it, is vital for understanding patterns of trait change in phytoplankton, and subsequent effects on ecology and biogeochemistry.

Phytoplankton diversity is often framed in terms of variation in phytoplankton functional traits, such as cell size (5,6). Functional traits of diatoms are well studied, and relationships between functional traits are usually considered as pairwise correlations, often involving fundamental tradeoffs between traits, such as cell size and nutrient uptake affinity (7,8). Based on this, trait-based models of diatoms and other phytoplankton often use the simplifying assumption that fitness is determined by a single master trait (e.g. size), and that all other traits are linked through a fixed correlation with the master trait (5). This therefore assumes that trait correlations are maintained during evolution, and that pairwise correlations are sufficient to describe constraints on complex phenotypes. However, multitrait relationships that extend beyond pairwise correlations can constrain phenotypic change on both plastic (9,10) and evolutionary (11) timescales. In addition, adaptive outcomes are consistent with these correlations shifting during adaptation (11). These multitrait relationships can be identified through statistical techniques such as principal component analyses (PCA), which can define the “trait-scape” for phytoplankton phenotypes (9) in one or more environments, for a set of traits that are commonly used to understand ecological or biogeochemical function. The trait-scape collapses multitrait variation onto a reduced set of axes, which allows us to study variation in multitrait phenotypes rather than in single traits. In contrast to master trait approaches, trait-scapes reveal shifts in trait correlations as well as in trait values, and automatically incorporate trait correlations beyond pairwise comparisons to include statistical correlations between three or more traits, whether or not knowledge of the biological basis for trait correlations is complete or correct. It is also worth noting that single locations in the trait-scape can define more than one multitrait phenotype (11), and that the single-trait measurements are useful for interpreting patterns of variation in, and movement on, trait-scapes.

Here, we used population bottlenecks to drive rapid diversification (12), which enabled us to explore patterns of evolutionary change in interrelated functional traits that affect the ecosystem and biogeochemical function of temperate diatoms. We did this both in a constant temperature environment and under moderate warming previously shown to drive adaptation in this genus even below the temperature optimum (13). Our motivation for using population bottlenecks was twofold. First, these conditions allowed diversification and fitness recovery in the absence of environmental change, which reveals how interrelated traits can vary through chance events and viability selection alone. Second, although chance events have the potential to contribute to diversification in populations of marine microbes, they are rarely used to generate diversity in experiments that subsequently allow adaptation to a changed environment. Chance events, such as drastic reductions in population size, have the potential to affect ecologically and biogeochemically important marine microbes. Diatoms, for example, can experience extreme fluctuations in population size both during bloom and bust growth, and as a result of the physical forces in ocean currents which can transport them relatively rapidly across different ecoregions (14,15). While there is ample evidence that natural selection does underly some of phytoplankton trait diversity, especially in terms of temperature niches (16–18), it is also likely that chance events play an important role. Specifically, populations adapting to temperature changes are often doing so either as they begin to bloom, or after a migration event, both of which involve chance acting immediately prior to selection, partially determining variation that natural selection has to act on.

We used repeated population bottlenecks to increase genetic and phenotypic divergence between replicate populations of diatoms. The experiment used six genetically and phenotypically distinct strains, in replicate (*see Fig. 1 for a schematic of the experiment*). For clarity, we refer to each starting genotype as a strain and to each evolving replicate population that is related by descent as a lineage. During bottlenecks, relaxed stabilizing selection allows new viable multitrait phenotypes to emerge. We refer to this as the reduced selection (RS) phase of the experiment. Many RS populations have lower population growth rates than their ancestors because chance, rather than natural selection, plays a dominant role in which cells survive. Using population bottlenecks to generate diversity is the basis of mutation accumulation experiments (19), which are commonly used in experimental evolution, mainly for the calculation of mutation rates and effect distributions. Most mutations accumulated during mutation accumulation experiments are deleterious, and we then observed how natural selection acts on this trait variation during fitness recovery by evolving the lineages as large populations (20). Natural selection is more effective in larger populations, so fitter multitrait phenotypes increase in frequency within each population during the full selection (FS) phase of the experiment. The experiment was ended when population growth rates stabilized. We took this to indicate (i) that FS populations were on or near local phenotypic optima, and that further adaptation would probably require an input of novel variation for fitness to eventually increase further (21), or (ii) that natural selection was acting on small differences in fitness, such that increases in fitness would be too slow to be relevant when considering single blooms or growing seasons.

**Figure 1.**
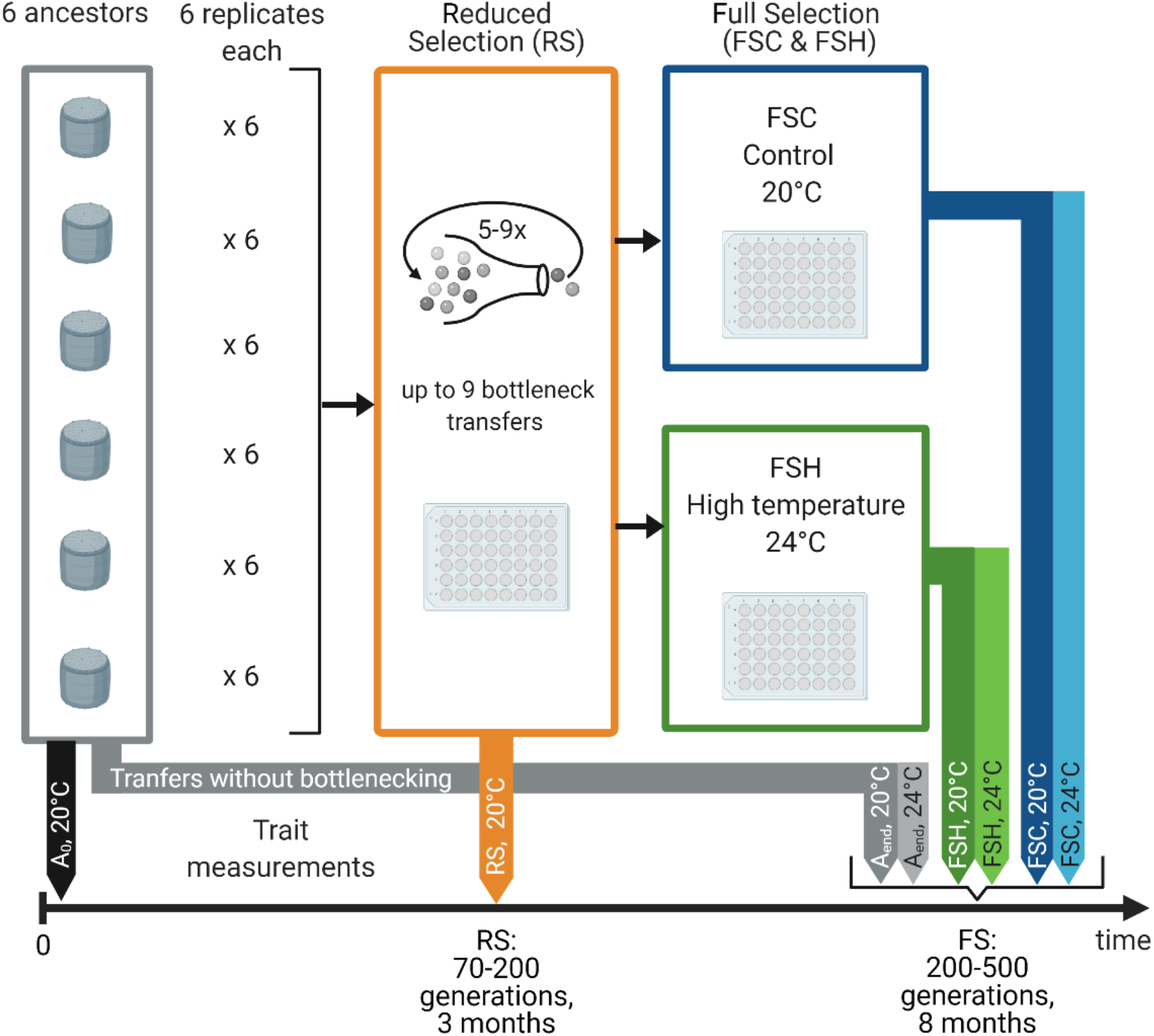
Schematic of the experimental design. Six strains (1010, 1050, 1587, 2929, and 3367) were used for the experiment. Six replicates from each strain (1010-1, −2, ..−6, 1050-1,…) underwent a procedure of up to 9 bottlenecks, each time transferring ~8 cells. After that, lineages were transferred regularly for 8 months under full selection with large inoculation sizes. The full selection was performed under control (1010-1C, 1010-2C,…) and high (+4 °C) temperature conditions (1010-1H, 1010-2H,…). Phenotypic trait measurements under control assay conditions took place at the beginning of the experiment (from the ancestral strains, A_0_), and at the end of the reduced selection phase (RS). At the end of the full selection phase, trait measurements of all ancestral (A_end_), evolved control (FSC) and high temperature lineages (FSH) were performed under control and high temperature assay conditions.

We consider whether populations returned to their ancestral multitrait phenotype during this fitness recovery (with or without environmentally-driven adaptation), moved to an area of the trait-scape occupied by a different genotype, thus connecting two peaks, or discovered a novel peak in the trait-scape (see Box 1). This study provides insight into patterns of multitrait variation in diatoms at multiple taxonomic levels, environments, and timescales. It also addresses the role of shifts in trait correlations during multitrait evolution, both in the absence and presence of environmental change, and contextualizes the role of evolution in trait correlations in terms of effects on growth rates and movement in trait-space.

### Box 1: Movement on trait-scapes

Movement in trait-scapes can be the result of changes in trait values, trait correlations, or both. Figure B1 shows possible types of movement of evolving populations on the trait-scape. Reduced selection will result in populations moving from their ancestral peak in the trait-scape (grey) to, on average, lower-fitness locations (orange). Movements in the trait-scape during the full selection phase are shown in blue. If trait values and correlations are strongly constrained, populations may recover trait values similar to their ancestor and be on “isolated” peaks. Alternatively, if trait values and correlations are less constrained, evolving populations may move to a new peak. If this new peak is in a previously occupied location (for example, by another genotype), then the two locations are “connected”. Otherwise, populations may move to a previously unoccupied location in the trait-scape under full selection, finding a “novel” peak. Understanding movement in trait-scapes can give insight into patterns of organismal change involving multiple traits values over multiple temporal and taxonomic scales.

**Figure B1.**
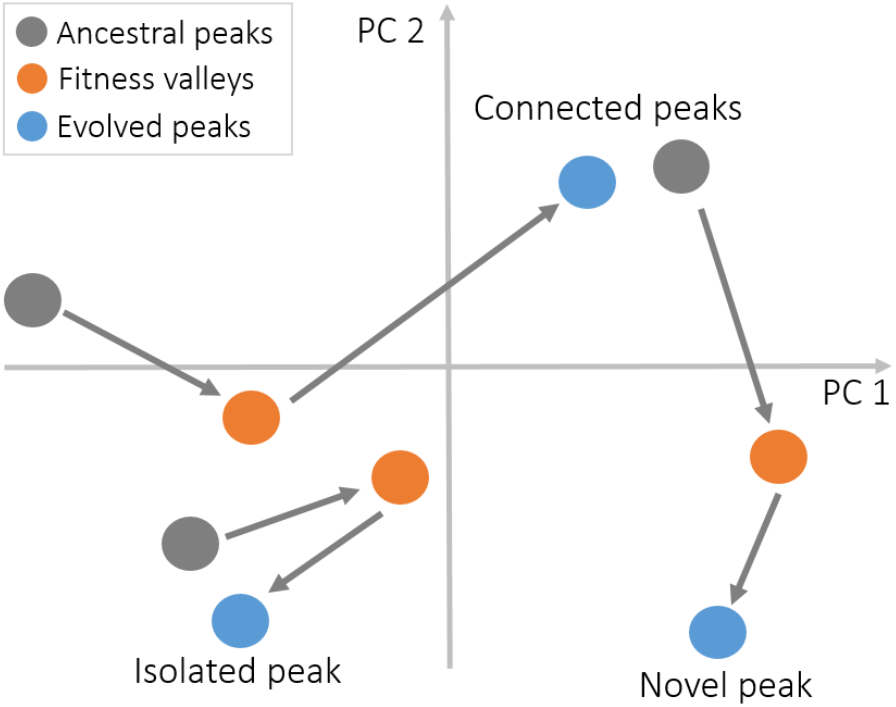
Schematic of the three possible movements in the trait-scape.

## Results

### Effectiveness of population bottlenecks and fitness recovery

The goal of our study was to examine patterns of variation generated in multitrait phenotypes in diatoms. We assume that our ancestral populations were well adapted to the culture conditions and thus sat on relatively high fitness peaks within the trait-scape. Our rationale for this assumption is that the ancestral strains had high enough fitness to be sampled and survive isolation, were all from culture collections, and had been grown in similar laboratory conditions for five months prior to the start of the experiment, during which time they had relatively high and stable population growth rates. The population bottlenecks resulted in strong fluctuations and an average reduction in population growth rate of 17-65% compared to ancestral rates (Fig. S3), and RS populations had different multitrait phenotypes than their ancestors (Fig. S6). Though growth rate may not be a complete description of fitness, large reductions in population growth rate in batch culture must have a negative impact on overall fitness. Following population bottlenecks, fitness recovery was effective, and full selection populations under control (FSC) and high temperature conditions (FSH) evolved higher population growth rates again that stabilized on average 20% below the growth of control populations propagated in the same environment as large populations (Fig. S4, F_1,143_ = 111, p<0.01)). Ancestral populations were maintained under control conditions as large populations and generally remained unchanged over the course of the experiment (*see methods*).

### The trait-scape

Previous work demonstrated that the ancestral multitrait *Thalassiosira* phenotypes could be studied using reduced axes (a trait-scape) that captured variation between genotypes (9,22). Here, we assess how variation generated within evolving single-genotype populations is captured by trait-scapes that also capture intra-genus variation.

Our experiments allowed us to assess how evolving populations move within the *Thalassiosira* trait-scape, focusing on whether evolution within lineages could produce new mulitrait phenotypes with new locations in the trait-scape, and whether different types of movements were seen in the trait-scape (*see Box 1*). Our rationale for exploring multitrait change using a trait-scape is that reduced axes have the potential to reveal correlations or tradeoffs that go beyond pairwise trait interactions, so that it simultaneously shows evolution in trait values and trait correlations. To do this, we combined the ancestral and evolved populations into one principal component analysis (Fig. 2). The first two principal components of the trait-scape explain almost 85% of the variation in trait values of nine functional traits (population growth rate, cell size, cell complexity, chlorophyll a content, particulate organic carbon and nitrogen, lipid content, silicic acid uptake, intracellular reactive oxygen; see Methods). Interspecific variation across strains (indicated by different colors) was significantly larger than within strain variation over the course of the experiment (F_1,11338_ = 1256, p<0.01), confirming that the trait-scape constructed from ancestral and evolved populations captured intraspecific variation. This was in good agreement with previous experiments using these ancestral strains (9). Most strains stayed on or returned to their ancestral location in the trait-scape after bottlenecks and subsequent evolution as large populations. Here, short periods of lineage evolution, such as would occur during individual blooms or growing seasons, do not usually overcome intragenus variation in these environments. Even with contributions from chance events and moderate warming, the majority of the evolving populations return to their ancestral location in the trait-scape when allowed to evolve as large populations. Overall, the trait-scape was evenly filled by the six strains, without large gaps that would indicate non-allowable trait value combinations within the space defined by the first two principal components. This result is consistent with previous experiments with these strains (9). Assessment of changes in single traits across the experiment and the movement of strains in the trait-scape including all reduced selection stages is provided in Fig. S6 and Fig. S7.

**Figure 2.**
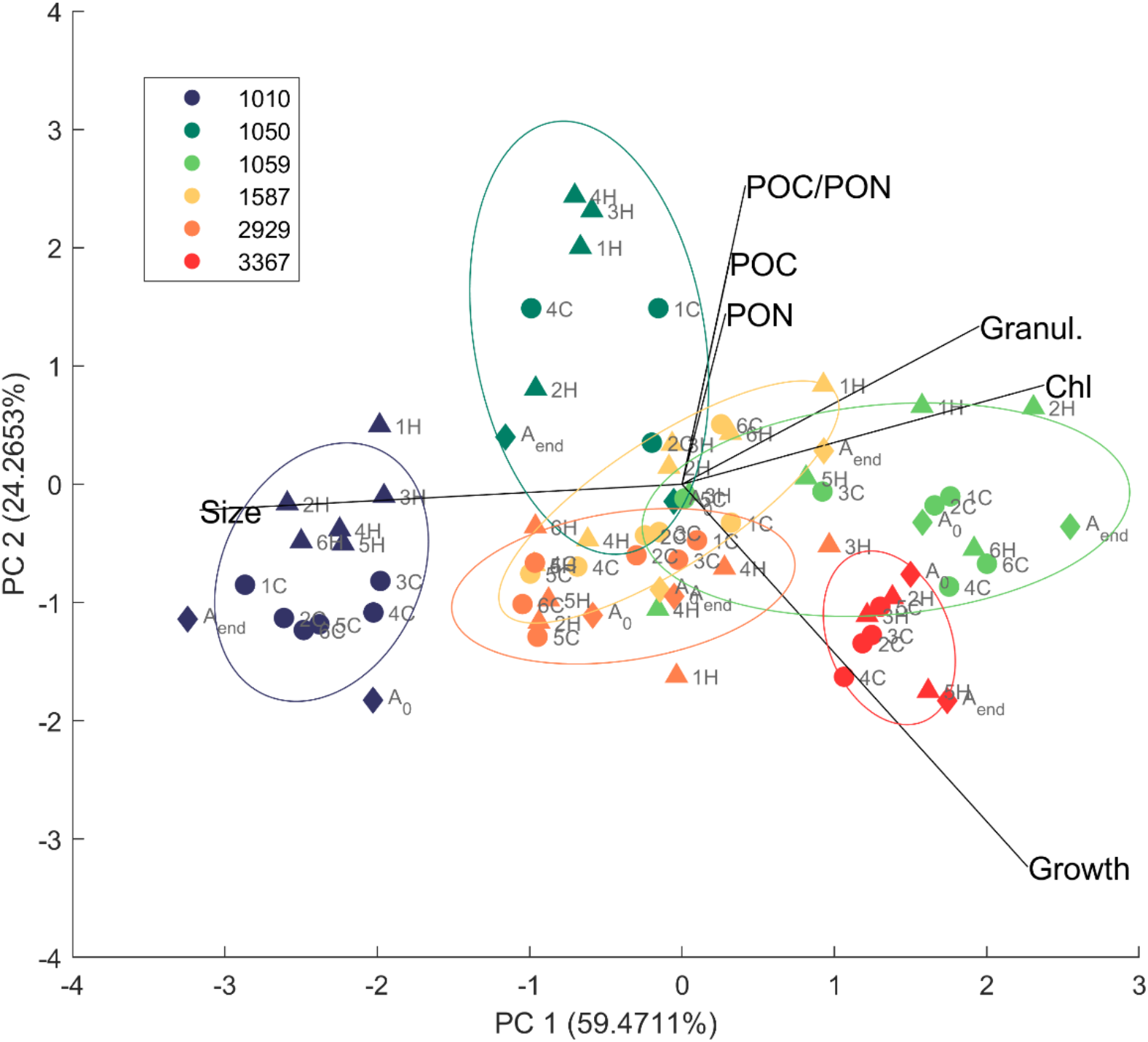
The trait-scape based on the principal component analysis of all ancestral (A_0_ and A_and_) and evolved (FSC and FSH) populations. Trait values were obtained under control conditions. Ancestors, FSC and FSH evolved populations are visualized together, colour-coded by strain. Ancestors are marked with a diamond, FSC populations with a circle, and FSH populations with a triangle. Replicates from the reduced selection are indicated by numbers, A_0_ and A_end_ mark the ancestral populations measured at the beginning and end of the experiment. Circles represent 75% confidence intervals for each strain.

### Movement of evolving populations on the trait-scape

We used the movement of populations on the trait-scape to define patterns of change to multitrait phenotypes in two environments. In the first, FSC populations evolved in the ancestral environment, so that any phenotypic variation between replicate populations of the same genotype are the result of population bottlenecks only. In the second, FSH populations evolved in a warmed environment, where environmentally-imposed selection could also contribute to movement on the trait-scape. We categorized movement in the trait-scape for each lineage as revealing either an isolated peak, a connected peak, or a novel peak (see Box 1, and “Analysis of trait-scape connectivity” in the Methods for more detail).

Despite most evolving populations returning to their ancestral location in the trait-scape, we observed all possible types of movement in the trait-scape (Box 1, Fig. 3), with evidence for isolated, connected, and novel peaks (summarized in Fig. 3d). For both FSC and FSH populations (62 populations total), nine of the 62 populations (14.5%) moved to a novel peak in the trait-scape, six of the 62 (6.7%) moved to a connected peak and 47 of the 62 (75.8%) returned to their ancestral peak. Exploration of the trait-scape was limited, but not absent, when populations evolved under full selection in the ancestral environment. Of the 31 FSC populations, only one population found a novel peak, and only two populations moved to a connected peak. The two populations that found connected peaks were reciprocal – a single pair of peaks was connected on this landscape during our experiment (1587-2929). This indicates that first, the trait-scape is reasonably well sampled in the ancestral environment and second, that it is possible for lineages to move on the trait-scape, and thus shift multitrait phenotypes, due to demographic perturbations alone.

**Figure 3.**
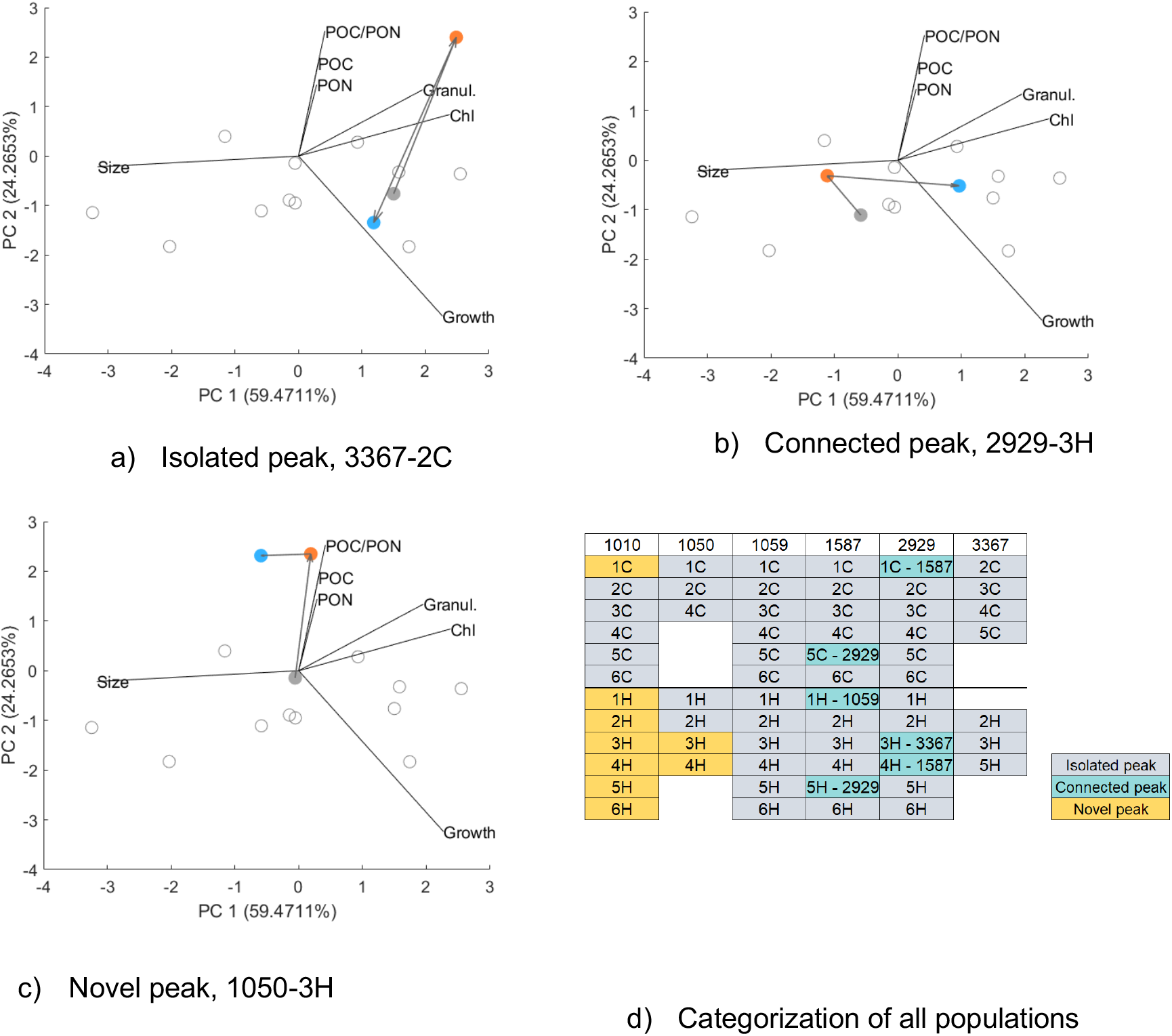
Example movements of three populations (3367-2C, 2929-3H, and 1050-3H) in the trait-scape (a-c). The principal component analysis was performed based on trait data from ancestral, FSC and FSH populations; reduced-selection populations were projected into this trait-scape. Data from all ancestors (A_0_ and A_end_) are depicted in grey, unfilled circles. Arrows follow the movements from an ancestral population (A_0_, grey filled circle) to its location after reduced selection (orange), and then to its location after full selection (blue). (d) Categorization of all populations into isolated, connected or novel peaks. In case of connected peaks, the connected strain is mentioned. Populations are sorted by strains, numbers represent reduced selection replicate identities, and letters indicate full selection conditions (C - control or H - high temperature).

Changes in multitrait phenotypes were more widespread when populations evolved in a different environment than their ancestor. Of the 31 FSH populations, four found connected peaks, evidencing three pairs of connected peaks on the trait-scape (1587-2929, 1587-1059, 3367-2929). In addition, eight FSH populations from two lineages located novel peaks, with strain 1010 often finding novel peaks. However, even when populations evolved in warmed environment, the ancestral locations in the trait-scape captured a large proportion of the locations occupied by the FSH populations - only two of the six strains located novel peaks, though all six strains evolved novel multitrait phenotypes (i.e. different values for some traits, see Fig. S7). This is surprising, given that warming is known to drive both plastic and evolutionary responses in *Thalassiosira*, so we expected more novel peaks (13,18). Our data suggests that the outcome of adaptation to moderate warming, including shifts in multitrait phenotypes, varies between strains, and between replicate populations (Fig. S7). However, in this experiment, 19 out of 31 FSH populations returned to their ancestral location in the trait-scape. Even though the multitrati phenotypes themselves changed, location in the trait-scape often stayed the same.

### Patterns of multitrait evolution

Population bottlenecks transiently relax the efficacy of natural selection, and one particularly interesting consequence of this for trait-based approaches is the potential for rapid shifts in trait correlations as well as trait values. While most of the evolving populations returned to their ancestral locations in the trait-scape, novel trait combinations did sometimes occur in FS populations.

Strains in locations categorized as isolated peaks in the trait-scape (1059 and 3367), showed no change from ancestral trait values when single traits are compared for the ancestral and FS populations of the same genotype (Fig. S7). Thus, even though many lineages evolved trait values different from their ancestors when selection was relaxed (RS phase) (Fig. S6, S7), most re-evolved ancestral trait values when selection was restored (FS phase). We were not able to measure all traits in the RS populations due to low growth and subsequent low biomass in these populations, so that changes in ROS, silica uptake and lipid content do not contribute to patterns on trait-scapes that include RS populations.

To categorize peaks using the full set of traits available in this study, we repeated the categorization above using only the ancestral and FS populations, so that the full set of traits could be used (Fig. 4, 5, S9). We found large increases in ROS in lineages 1050-3H and 1050-4H, suggesting that our categorization of movement based on a trait-scape that includes the RS populations, where ROS was not measurable, may underestimate novel peaks.

**Figure 4.**
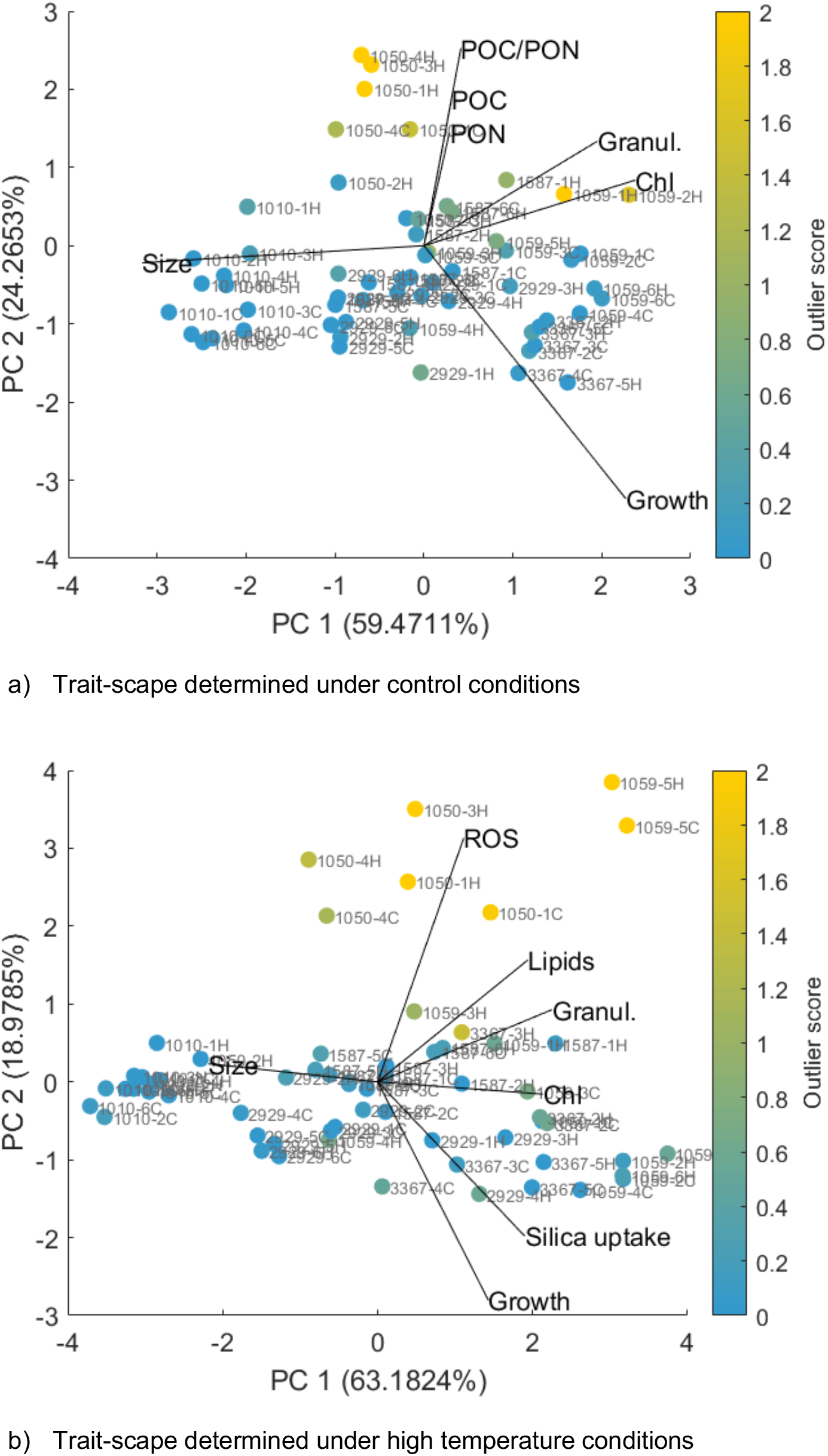
The trait-scape of all FSC and FSH population measured under control (a) and high temperature (b) conditions, with outliers from ancestral trait correlations indicated by colour. Traits included in the principal component analysis differed between the two temperature conditions to include as many traits as possible, whilst maintaining comparability to the trait-scape that was calculated based on the available trait data of all ancestral, reduced selection and evolved lineages.

**Figure 5.**
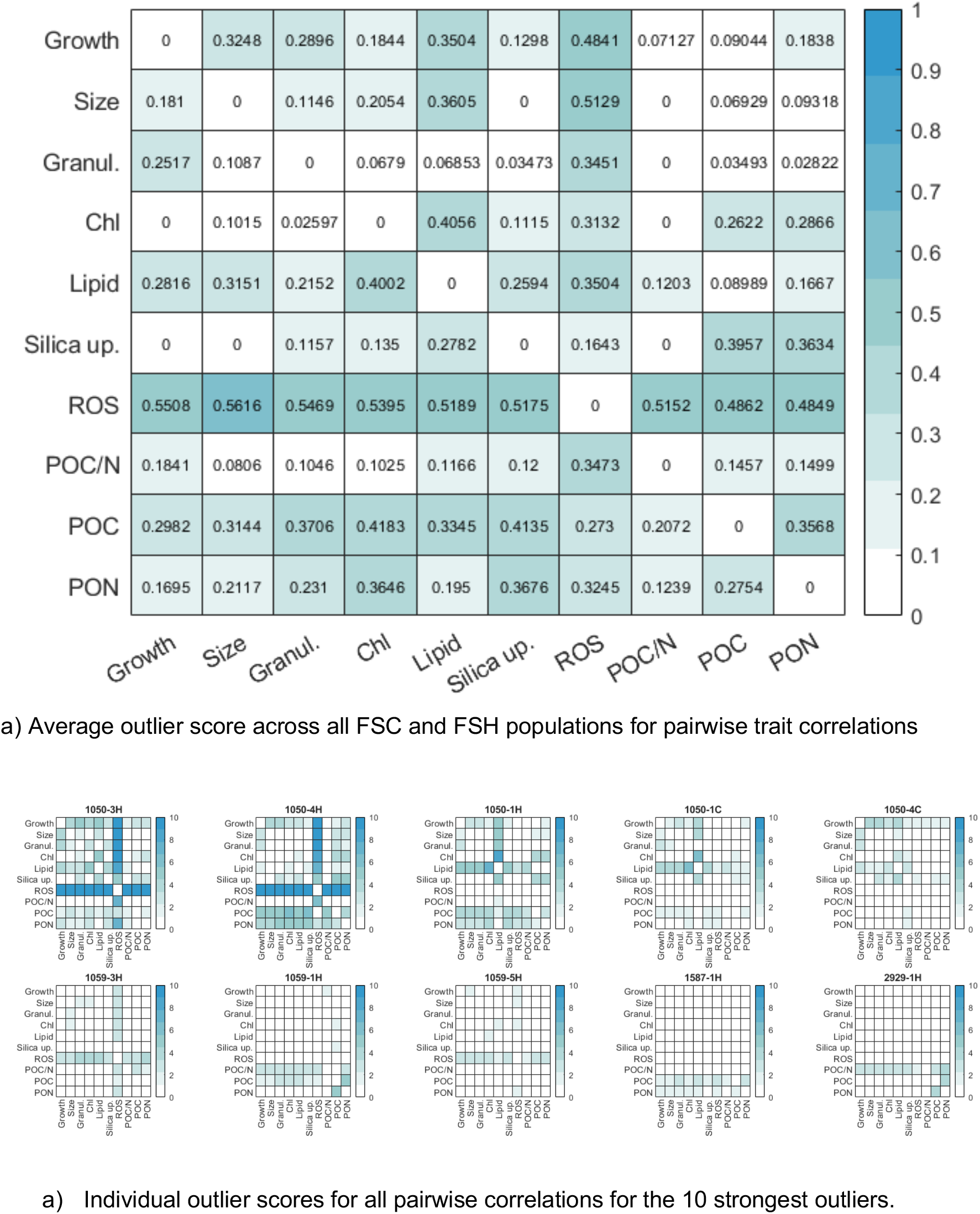
Outlier analysis for all pairwise trait correlations across strains (a) and for the 10 strongest outliers (b).

Interestingly, populations moving between connected peaks in the trait-scape did not evolve the same set of trait values as the ancestral strain originally on that peak (Fig. S11, S12). Instead, they developed multitrait phenotypes distinct from both their own ancestor and the ancestors or FS populations of other strains. This agrees with simulations of adaptation on trait-scapes (11) that predict the evolution of cryptic phenotypes, where more than one multitrait phenotype exists in a single location in trait-scape. This is exciting, because while novel multitrait phenotypes evolved, trait evolution was captured within the pre-existing trait-scape, and often involved moving to a location in the trait-scape occupied by a different strain. The number of possible multitrait phenotypes present in a single location in the trait-scape could be reduced by including more traits that load on the first two axes of the trait-scape. Candidates for additional traits are those that are likely to vary both between species, and between replicate populations evolving in changed environments. In the case of changing temperature, respiration measurements are one such candidate trait (23).

### The role of trait correlations in the evolution of novel phenotypes

First, we examined whether novel multitrait correlations were associated with particular trait values. To do this, we quantified an outlier score for each FS population to quantify shifts in trait correlations. This score identifies populations where the relationship between two traits has shifted substantially from the ancestral correlation (see ”Outlier analysis” in the Methods section). We found that some trait values were indeed associated with high outlier scores. For FSC populations that evolved in the ancestral environment, we found that populations with high particulate organic carbon (POC) and nitrogen (PON) content, as well as high granularity and chlorophyll content depart most from ancestral trait correlations, whereas populations with large cell sizes and high growth rates depart least from ancestral trait correlations (Fig. 4(a)). FSH populations that evolved in the warmed environment showed a similar pattern, where populations with large cell sizes and high growth rates had lower outlier scores (Fig. 4(b)). In FSH populations, where temperature was elevated by 4°C, many populations with high outlier scores also had high reactive oxygen species (ROS) values, which suggests that large departures from ancestral trait correlations may be stressful for the cells (Fig. S9). This is consistent with previous work showing that ROS production is a common response to stressful environmental conditions (24–26).

The above examined how particular trait values are associated with higher outlier scores. A related question is whether specific traits, and associated trait correlations, are more likely to evolve than others. To answer this, we compared the mean residual values across strains for each pairwise trait correlation (Fig. 5(a)). Nine out of the ten most extreme outliers evolved under high temperature, indicating that evolution in a new environment may favor changes in trait correlations (Fig. 5(b)). Across all outliers, the largest outlier scores involved correlations with ROS, POC, PON, and lipid content (Fig. 5(a)), although more modest departures from ancestral correlations occurred in many other pairwise trait correlations. Taken together, this indicates that high outlier scores are typically associated with a substantial and directionally consistent (i.e. increase or decrease) change in a few key traits. This means that some traits are more likely to be associated with departures from ancestral trait correlations than are others. Second, it shows that some trait correlations evolve more readily than others, even without environmental change.

While trait correlations evolved in most populations, the identity of the trait correlations that departed most from ancestral values were population-specific (Fig. 5(b)). 1050-3H, 1050-4H, 1059-3H and 1059-5H showed the most change from the ancestral correlations between ROS and other traits. 1050-1C and 1050-1H diverged most from ancestral correlations between lipid content and other traits, and 1059-1H and 1587-1H showed the most change from ancestral correlations between POC and PON content. Strains 1050 and 1059 had the greatest number of outlier populations, suggesting that these strains have more flexibility in trait correlations than the other strains in this study. Overall, the strongest departure from ancestral trait correlations was related to large shifts in correlation due to changes in a single trait (typically ROS).

Considering the relationship between growth and outlier scores provides information about the potential fitness effects of evolving new trait correlations, as population growth impacts fitness, especially during diatom blooms. We found that populations with large changes in trait correlations had up to 60% reduction in growth rates relative to their ancestral strain (Fig. S10, F_1,55_ = 30.662, p<0.01), even though these growth rates had stabilized by the end of the experiment. When these populations were then grown for additional time (10 months with weekly transfers), growth rate did increase further, ranging between ±20% of ancestral growth. This indicates while trait correlations evolve readily after population bottlenecks, new trait correlations are likely to have a detrimental impact on fitness. These correlations may then change further or revert if there is the opportunity for continued adaptation under conditions that select on rapid growth. It is worth noting, however, that the increase in growth rate occurred after long-term culturing under conditions that consistently favored rapid population growth (batch culture). In contrast, typical diatom blooms may last weeks, and growth seasons for temperate diatoms typically span a few months. Taken together, our data show that shifts in trait correlations, even if they are transient, have the potential to play a role in determining evolutionary trajectories, as well as underly variation in multitrait phenotypes expressed early in adaptation.

## Discussion

Observed variation in diatom phenotypes is generated by a variety of evolutionary forces including natural selection and chance events. The relationships between the many traits that define these phenotypes can be assessed using a trait-scape. Here, we used an evolution experiment where six phenotypically distinct strains of the diatom genus *Thalassiosira* were diversified using repeated population bottlenecks and were subsequently re-evolved as large populations. We demonstrate that novel phenotypes emerged even in the absence of environmental change and that the phenotypic shifts in the evolved populations could be studied using a trait-scape defined by the ancestral populations. This suggests two fundamental insights. First, chance events such as repeated population bottlenecks can provide phenotypic variation on which natural selection subsequently acts, whether or not the environment changes. This is likely to be important given the bloom and bust growth strategies of marine diatoms and the dynamic environments in which they live; the nature of this variation fixed by chance events represents part of the variation that natural selection subsequently acts on. Understanding the possible routes that environmentally-driven evolution can take depends on which phenotypic starting points are available for selection to act on. We show that in most cases, the bottlenecked populations simply revert to their ancestral multitrait phenotypes when subsequently evolved as large populations, but different evolutionary starting points provided by bottlenecks do occasionally allow novel multitrait phenotypes to evolve in large populations, even in a constant environment. The growth strategies of marine microbes coupled with their dynamic ocean habitats point to frequent bottlenecking in natural populations. While a low proportion of bottlenecked populations evolved new multitrait phenotypes, when this happens repeatedly over many populations, as will be the case in the ocean, population bottlenecks could generate substantial diversity in diatom functional traits and their correlations. This is consistent with bottlenecks generating diversity in other taxa that experience large fluctuations in population size, such as viruses and bacteria (27,28).

The second insight is that multitrait variation is constrained to few fixed axes of variation where all of the functional traits used in this study projected strongly. This indicates that considering changes to multitrait phenotypes made up of numerous intertwined functional traits can reveal patterns of phenotypic evolution, at least in the absence of dramatic or stressful environmental change. This is useful because it does not depend on knowledge of how traits are linked to one or more other traits. This may be especially relevant for ecosystem modelling, where phytoplankton multitrait phenotypes are often simplified using a master trait, which determines the values of all other functional traits via fixed trait correlations (5,6). Here we demonstrate that, for our system, there is no master trait that can be used to understand how multitrait phenotypes evolved – all of the traits projected strongly onto the two trait-scape axes. Instead, repeatable multitrait evolution occurs along a major axis of variation defined by several common diatom functional traits and the relationships among them. Because these functional traits are relevant to other phytoplankton functional groups (29), we expect the concept, utility, and many of the relationships revealed in any given phytoplankton trait-scape to generalize to other phytoplankton, especially within functional groups (silicifiers, calcifiers). This is because all phytoplankton have similar functional traits, and many are linked to basic, conserved metabolism (30). We do not expect that all functional groups have exactly the same trait-scape, as functional groups with different biogeochemical functions and environmental niches will most likely have different trait axes. However, trait-scapes can provide a reduced dimensional way to assess multitrait phenotypic changes. Trait-scapes are also a useful because they provide a way to capture evolution in trait correlations. Our findings suggest that master-trait approaches that rely on fixed correlations are inappropriate for studying multitrait change at this taxonomic level and timescale in diatoms.

Our study used two environments, both of which were nutrient replete and below the temperature optimum for these strains. While a similar degree of warming has been shown to drive adaptation in large laboratory populations of diatoms (17,31), major shifts in fundamental growth strategies, such as those associated with heat stress or nutrient starvation, would not be expected. While our FSH populations did not experience heat stress, an increase of 4°C is in line with the magnitude of environmental change expected in many marine systems over the next 100 years (32). This is also the magnitude of change that a phytoplankton population might experience due to mixing between adjacent water masses (33), or growing sequentially over several seasons (34).

While individual traits changed in response to warming, multitrait phenotypes often returned to the ancestral location in the trait-scape. Based on this, we hypothesize that relative to individual trait values, multitrait phenotypes may be more stable across environmental gradients that do not provoke major shifts in growth strategies. Thus, trait-scapes can offer insight into fundamental constraints on functional trait variation within phytoplankton groups. However, the stability of the trait-scape in the face of novel or stressful environments is unknown, and should be explored in future studies.

## Conclusions

Phytoplankton play crucial roles in aquatic food webs and biogeochemical cycling, and they do so through their interconnected web of functional traits. The trait-scape described by functional trait values and correlations in our study is robust to variation produced in the short term by chance events and natural selection, and across two similar and non-stressful environments. Most populations return to their ancestral locations in the trait-scape after demographic perturbations. However, a minority of populations evolve different multitrait phenotypes from their own ancestors. These new phenotypes can be captured by movement on the trait-scape defined by the ancestral populations, often moving to areas of the trait scape that are already occupied (connecting peaks). Trait-scapes offer an alternative to master-trait based views of how trait correlations constrain multitrait phenotypes in diatoms, and we suggest that this same approach will also be useful in other phytoplankton functional groups. This multi-trait framework incorporates higher-level trait relationships and is a tractable way to understand and represent trait-based variation in phytoplankton.

## Materials and Methods

### Diatom cultures

Six strains of *Thalassiosira sp*. from the National Centre for Marine Algae and Microbiota at Bigelow were used: CCMP 1010, 1050, 1059, 1587, 2929 and 3367 (Table S1). These differed from each other in original isolation location, as well as traits such as cell size and population growth rate (see ref. (9) for a detailed description of the strains). Strains were chosen that did not clump or form chains in our culture conditions to allow easy isolation of a few cells for bottleneck transfers. Data from previous experiments using 13 different *Thalassiosira* strains including these six, confirmed that the strains used in this study are a representative divergent sample of a larger collection of genotypes (9).

### Culture maintenance

Cultures were grown in sterile f/2 media (35) made from natural sea water (collected in St Abbs, United Kingdom), at 20°C and approximately 60 μmol photons m^-2^ s^-1^ (measured with a 4-pi sensor) at a 12h:12h light:dark cycle. For the evolution experiment, cultures were maintained in transparent 48-well plates covered with Breath-Easy breathable plate-seals (Sigma Aldrich). Ancestral populations were maintained under identical nutrient, light and temperature conditions, but in 40 mL culture flasks. Cultures were diluted every 7-10 days with fresh media to transfer sizes between 1000-2000 cells/ml to maintain exponential growth. These culture conditions are the control conditions.

### Evolution experiment

The experiment was divided into two phases, an initial 3-month long reduced-selection (RS) phase (corresponding to 70-200 generations) followed by an 8-month full selection (FS) phase (200-500 generations) (Fig. 1).

The reduced-selection phase of the experiment was initiated with six replicates of each of the six experimental strains, resulting in a total of 36 experimental populations. Due to the extinction of three populations within the reduced-selection phase, 33 populations entered the full selection phase of the experiment. The full selection was performed in a control (FSC, 20°C) and a warm environment (FSH, 24°C). The warmed environment is below Topt for these strains and is not an obviously stressful growth environment.

Trait measurements of populations were performed at the beginning of the experiment, at the end of the reduced-selection phase and at the end of the full selection phase. During the experiment trait values were measured under control conditions, at the end of the experiment, trait values were obtained under control (20°C) and high temperature (24°C) assay conditions. Table S2 lists which trait values were collected for each experimental stage. Analyses have a focus on the data collected under control conditions. Whenever data from high temperature assay conditions is analysed, this is stated.

#### Reduced-selection phase

To reduce the efficacy of natural selection and allow divergent lower-fitness phenotypes to evolve, we repeatedly induced population bottlenecks (~8 cells) in the 36 experimental populations (six replicates of six ancestral strains). Initially, bottlenecks were induced every 7 days (corresponding to an average of 13 generations). As growth rates decreased over time, this time window was increased to every 14 days towards the end of the experiment (corresponding to an average of 18 generations). For laboratory feasibility, all populations were bottlenecked at the same time, as long as they had reached a minimum cell concentration of 2000 cells/mL. If cell concentrations were lower than 2000 cells/mL, cultures were diluted to 500 cells/mL instead, to allow for population recovery before a new bottleneck was induced.

The bottlenecking was performed in two dilution steps. First the most recent inoculation of each population was diluted to 2000 cells/mL. From this dilution the actual bottlenecking was performed aimed to transfer ~8 cells into 1 mL fresh media.

The bottlenecking was repeated 5-9 times, depending on the lineage. This corresponds to a total experiment length of 3 months and 70-200 generations, depending on population growth rates. Especially towards the end of the bottlenecking phase, single populations had decreased in growth to such a degree that they did not grow. In these cases the previous transfer was used to induce a new bottleneck. Throughout the reduced-selection phase the maximum growth rate was monitored via *in-vivo* fluorescence. At the end of the bottleneck phase, population growth rates were reduced by an average of 45% compared to ancestral growth rates (Fig. S3).

#### Full selection phase

During the subsequent full selection phase populations were propagated by batch culture with transfer sizes of ~1000 cells every 7 days. 33 populations entered this phase. The full selection was performed in two environments, a control (20°C) and a warm environment (24°C) to allow for the later comparison of plastic and evolutionary responses to different conditions. The warm environment was chosen to create a selection regime that clearly diverges from the control environment without stressing the weakened reduced-selection populations to such a degree that they would die (based on the provided information on temperature ranges for the ancestral populations by the Bigelow culture collection). As an example, from the ancestral strain 1010, 6 replicates underwent the reduced selection (called 1010-1 to 1010-6), in the full selection phase each replicate was exposed to two temperature environments (named 1010-1C to 1010-6C and 1010-1H to 1010-6H). In total, populations were transferred ~25 times in the full selection phase of the experiment, corresponding to 200-500 generations. Maximum growth rates were measured every 5 to 10 transfers throughout the full selection phase to monitor fitness recovery. Before the final trait measurements the growth rate of all populations was measured over four transfers (four weeks) to ensure that the population growth rates had stabilized, indicating that populations were on or near a local high-fitness solution (Fig. S4, S5).

### Trait measurements

To investigate how trait values changed throughout the experiment, trait measurements were performed at the onset of the experiment (A0), at the end of the reduced-selection phase (RS) and at the end of the full selection phase (FSC, FSH). At the end of the full selection phase, the experimental ancestors, which were maintained under control conditions throughout the experiment, were measured again to account for any trait change during normal culture maintenance (Aend). Where applicable, all traits were measured during mid-exponential growth phase and daily mid-light phase. A detailed illustration of single trait changes across the experiment can be found in Fig. S7.

Trait measurements at the beginning and at the end of the reduced-selection phase included maximum growth rate, cell size, cell complexity, chlorophyll a content, and particulate organic carbon and nitrogen content. At the end of the full selection phase, a more extensive set of traits were measured for the FSC- and FSH-populations and the experimental ancestors, in both the control and the hot environment, Table S2. Pilot data showed that populations reach stable trait values after acclimating for 7 days to the test environment (9,22).Therefore, trait measurements were performed for all populations after an acclimation period of 7 days (one transfer) in the test environment. All trait measurements are described below.

#### Growth rate

Growth rates were measured via daily *in-vivo* fluorescence using a plate reader (excitation: 455nm, emission: 620nm; Tecan Spark, United Kingdom). The exponential growth rate was calculated for each time step using:

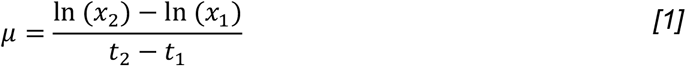

Maximum growth rates were determined based on the average growth over 4 consecutive time steps. During the bottleneck phase growth rates were determined based on single replicates per evolving population; for the final trait measurements growth rates of all ancestral and back-selected populations were determined based on 3 replicates per evolving population.

#### Cell size, cell complexity and chlorophyll a

Cell size, cell complexity and chlorophyll a concentration of all populations were determined by flow cytometry. Cell size estimates were determined based on the median forward scatter area, and converted to μm using a standard curve based on standard beads of 1, 3, 6, 10 and 25 μm bead size. The resulting values for cell size corresponds to the cell diameter.

Cell complexity was estimated based on median side scatter area, chlorophyll a (Chl a) was determined based on median fluorescence in the PerCP-Cy5-5-A channel (488nm wavelength, filter 695/40BP).

#### Particulate organic carbon and nitrogen

Particulate organic carbon (POC) and nitrogen (PON) content were determined using elemental analysis. Populations were grown in 50 mL culture flasks containing 45 mL of media, up to a cell density of ~112,000 cells/mL. The culture was split into two technical replicates of 20 mL, which were filtered onto pre-combusted Whatman GFF filters. From the remaining sample volume cell concentrations were determined using *in-vivo* fluorescence and previously determined fluorescence standard curves for each strain. The dry weight of the cells was determined after drying the filters over 2 days at 40°C. Filters were then treated to remove potential inorganic carbon. To do so, the filters were dampened with 200 μL distilled water, and 100 μL 2M hydrochloric acid was then applied to remove any inorganic carbon. After air-drying filters overnight, the samples were analyzed in a Thermo Fisher Scientific FlashSMART 2000 Elemental Analyzer.

#### Lipid content, silicic acid uptake and production of reactive oxygen species

The neutral lipid content, silicic acid uptake and production of reactive oxygen species were determined based on (22). In summary, neutral lipid content of cells was determined by adding a BODIPY 505/515 stain to the samples, at a final concentration of 2.6 μg/mL of BODIPY. After 10 min incubation, samples were analysed in the flow cytometer using median FITC-A (488nm wavelength, filter 530/30 BP).

Silicic acid uptake was calculated from PDMPO uptake by inoculating 500 μL samples with 0.125 μmol/L PDMPO dye (LysoSensor™ Yellow/Blue DND-160, Invitrogen) for 24 h, blank samples were inoculated in the same way excluding the dye. PDMPO is incorporated by cells with the same rate as silicic acid, allowing an estimate of the rate of silica production. After incubation under 12h:12h light:dark conditions silicic acid uptake was determined using median AmCyan-A of the flow cytometer (405nm wavelength, filter 525/50 BP), subtracting values of undyed samples from the dyed samples.

Relative intracellular ROS as a measure of stress was determined using H2DCFDA dye. 500 μL sample volumes were inoculated for 5 hours in the dark in the test environment (20/24°C) with a concentration of 80 μmol/L dye. Blank population samples excluding dye were inoculated in the same manner. After incubation relative intracellular ROS was calculated from H2DCFDA fluorescence using a Tecan Spark plate reader (excitation: 488nm, emission: 525nm) and blank values were subtracted from the dyed sample values.

### Data analysis

Data were analysed and plotted using Matlab, statistical analyses were performed in the R environment.

#### Correction for influence of cell size on other traits

Cell size is known to be a major functional trait, influencing many other functional traits; many metabolic traits show a linear correlation with cell size (36). We corrected the analyses of correlation matrices and principal components by the influence of cell volume. Traits that were corrected for cell volume were granularity, Chl a, lipid content, ROS production, silicic acid uptake, POC content and PON content. Growth rate and POC/PON ratio were not corrected for cell volume, since there is no linear relationship between growth rate and cell size (36) and since the ratio between POC and PON should not necessarily be correlated with cell size. The cell volume correction was performed by dividing the trait value of each sample by the corresponding calculated cell volume. Cell volume was calculated based cell diameter, assuming a spherical shape. Since only the diameter but not the height of the cylindrical diatoms were known, we calculated the spherical volume.

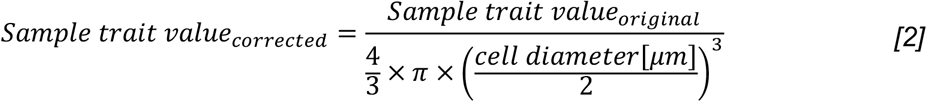

#### Principal component analysis

Principal component analyses (PCA) were performed after standardizing all trait data. The number of traits that were included in individual analyses differed slightly from each other, since not all traits were measured in all environments for all experimental stages (Table S2; see also section above on trait measurements for details). To project other populations into a defined PCA space, their normalized trait values are multiplied by the coefficients of the PCA space.

#### Analysis of trait-scape connectivity

To analyse the connectivity of the trait-scape, high-fitness (ancestral, FSC and FSH) locations were categorized as isolated, connected or novel. To maximize the comparability between analyses, this analysis was performed based on the PCA including trait values of ancestral and full selection populations under control assay conditions. Trait values of reduced selection populations were projected into the trait-scape.

First, populations that returned to the ancestral peak were identified by calculating the Euclidean distance between ancestral peak and location after reduced selection, and the distance between ancestral peak and location after full selection. If the latter distance was smaller than the former, the peak of the full selection population was classified as an isolated fitness peak.

For all already classified isolated fitness peaks, a 75% confidence interval was calculated around the ancestral peak. All populations that did not meet the first criterion, but remained in this 75% confidence interval of their ancestor were also classified as isolated fitness peaks. Any population that was not classified as an isolated fitness peak and that moved into the 75% confidence interval of another strain, was classified as a connected fitness peak.

All remaining populations that were neither classified as having an isolated peak nor a connected fitness peak, were considered to have found a previously unoccupied location in trait-scape, referred to as novel fitness peak.

#### Outlier analysis

To identify outliers among the FSC and FSH populations that diverged from ancestral trait values and trait correlations, all possible pairwise trait correlations were calculated for ancestors, FSC and FSH populations under control conditions. Across all ancestral strains linear regressions with a 95% confidence interval were calculated (Fig. S8). The residual values for FSC and FSH populations that diverged from the ancestral 95% confidence were calculated, all residual values for trait correlations that stayed within the ancestral confidence interval were set to zero. For each population an outlier score was calculated as the mean residual value across all pairwise trait correlations.

#### Statistical analysis

To compare the intra-vs. interspecific variation in trait-scape, Euclidean distances between all ancestral, FSC and FSH populations were calculated based on their location in the trait-scape using Matlab. The resulting dataset was classified by strain and grouped into intraspecific or interspecific Euclidean distances. Statistical analysis was then performed in the R environment using the nlme and lme4 packages. Since both a visual check via a Q-Q plot and the Levene test suggested that the data is not normally distributed, a generalized linear mixed-effects model was applied with type of variation (inter-vs. intraspecific) as fixed effect and strain as random effect. The effect of bottlenecking and backselection on growth rates was analyzed using a a generalized linear mixed-effects model with the experimental stage, respectively final outlier score as fixed effect strain as random effect.

## Data accessibility

All trait data can be found in the supplementary material for this manuscript. Code used for analysis will be deposited on Github upon paper acceptance. During review, code can be found at https://bit.ly/CodesHinnersetal

## Acknowledgments

The authors would like to thank Toby Samuels, Carmen Robinson and Katy McDonald for help with lab work. Flow cytometry data were generated within the Flow Cytometry and Cell Sorting Facility in the School of Biological Sciences at the University of Edinburgh. Support for flow cytometry was provided by Martin Waterfall. Particulate organic carbon and nitrogen analyses were performed by the Chemical Analysis Facilities of the School of Geosciences at the University of Edinburgh, with support from Gavin Sim and Joe Casillo.

## References

1. Lande R. Natural selection and random genetic drift in phenotypic evolution. Evolution. 1976;314–34.

2. Henson SA, Cael B, Allen SR, Dutkiewicz S. Future phytoplankton diversity in a changing climate. Nat Commun. 2021;12(1):1–8.

3. Sjöqvist C, Godhe A, Jonsson P, Sundqvist L, Kremp A. Local adaptation and oceanographic connectivity patterns explain genetic differentiation of a marine diatom across the North Sea–Baltic Sea salinity gradient. Mol Ecol. 2015;24(11):2871–85.

4. Rengefors K, Kremp A, Reusch TB, Wood AM. Genetic diversity and evolution in eukaryotic phytoplankton: revelations from population genetic studies. J Plankton Res. 2017;39(2):165–79.

5. Ward BA, Dutkiewicz S, Jahn O, Follows MJ. A size-structured food-web model for the global ocean. Limnol Ocean. 2012;57(6):1877–91.

6. Dutkiewicz S, Cermeno P, Jahn O, Follows MJ, Hickman AE, Taniguchi DA, et al. Dimensions of marine phytoplankton diversity. Biogeosciences. 2020;17(3):609–34.

7. Litchman E. Trait-Based Diatom Ecology. In: The Molecular Life of Diatoms. Springer; 2022. p. 3–27.

8. Lindemann C, Fiksen Ø, Andersen KH, Aksnes DL. Scaling laws in phytoplankton nutrient uptake affinity. Front Mar Sci. 2016;3:26.

9. Argyle PA, Walworth NG, Hinners J, Collins S, Levine NM, Doblin MA. Multivariate trait analysis reveals diatom plasticity constrained to a reduced set of biological axes. ISME Comm. 2021;1(1):1–11.

10. Malcom JW, Hernandez KM, Likos R, Wayne T, Leibold MA, Juenger TE. Extensive cross-environment fitness variation lies along few axes of genetic variation in the model alga, C hlamydomonas reinhardtii. New Phytol. 2015;205(2):841–51.

11. Walworth NG, Hinners J, Argyle PA, Leles SG, Doblin MA, Collins S, et al. The evolution of trait correlations constrains phenotypic adaptation to high CO2 in a eukaryotic alga. Proc R Soc B Biol Sci. 2021;288(1953):20210940.

12. Naciri Y, Linder HP. The genetics of evolutionary radiations. Biol Rev. 2020;95(4):1055–72.

13. Schaum CE. Enhanced biofilm formation aids adaptation to extreme warming and environmental instability in the diatom Thalassiosira pseudonana and its associated bacteria. Limnol Ocean. 2019;64(2):441–60.

14. Doblin MA, Van Sebille E. Drift in ocean currents impacts intergenerational microbial exposure to temperature. Proc Natl Acad Sci USA. 2016;113(20):5700–5.

15. Jönsson BF, Watson JR. The timescales of global surface-ocean connectivity. Nat Commun. 2016;7(1):1–6.

16. Irwin AJ, Finkel ZV, Müller-Karger FE, Ghinaglia LT. Phytoplankton adapt to changing ocean environments. Proc Natl Acad Sci. 2015;112(18):5762–6.

17. O’Donnell DR, Hamman CR, Johnson EC, Kremer CT, Klausmeier CA, Litchman E. Rapid thermal adaptation in a marine diatom reveals constraints and trade-offs. Glob Change Biol. 2018;24(10):4554–65.

18. Jin P, Agustí S. Fast adaptation of tropical diatoms to increased warming with trade-offs. Sci Rep. 2018;8(1):1–10.

19. Halligan DL, Keightley PD, others. Spontaneous mutation accumulation studies in evolutionary genetics. Annu Rev Ecol Evol Syst. 2009;40:151.

20. Estes S, Lynch M. Rapid fitness recovery in mutationally degraded lines of Caenorhabditis elegans. Evolution. 2003;57(5):1022–30.

21. Lenski RE, Wiser MJ, Ribeck N, Blount ZD, Nahum JR, Morris JJ, et al. Sustained fitness gains and variability in fitness trajectories in the long-term evolution experiment with Escherichia coli. Proc R Soc B Biol Sci. 2015;282(1821):20152292.

22. Argyle PA, Hinners J, Walworth NG, Collins S, Levine NM, Doblin MA. A High-Throughput Assay for Quantifying Phenotypic Traits of Microalgae. Front Microbiol. 2021;2910.

23. Padfield D, Yvon-Durocher G, Buckling A, Jennings S, Yvon-Durocher G. Rapid evolution of metabolic traits explains thermal adaptation in phytoplankton. Ecol Lett. 2016;19(2):133–42.

24. Malerba ME, Marshall DJ. Testing the drivers of the temperature–size covariance using artificial selection. Evolution. 2020;74(1):169–78.

25. Carrara F, Sengupta A, Behrendt L, Vardi A, Stocker R. Bistability in oxidative stress response determines the migration behavior of phytoplankton in turbulence. Proc Natl Acad Sci USA. 2021;118(5):e2005944118.

26. D’Autréaux B, Toledano MB. ROS as signalling molecules: mechanisms that generate specificity in ROS homeostasis. Nat Rev Mol Cell Biol. 2007;8(10):813–24.

27. Windels EM, Fox R, Yerramsetty K, Krouse K, Wenseleers T, Swinnen J, et al. Population bottlenecks strongly affect the evolutionary dynamics of antibiotic persistence. Mol Biol Evol. 2021;38(8):3345–57.

28. Bull RA, Luciani F, McElroy K, Gaudieri S, Pham ST, Chopra A, et al. Sequential bottlenecks drive viral evolution in early acute hepatitis C virus infection. PLoS Pathog. 2011;7(9):e1002243.

29. Litchman E, de Tezanos Pinto P, Edwards KF, Klausmeier CA, Kremer CT, Thomas MK. Global biogeochemical impacts of phytoplankton: a trait-based perspective. J Ecol. 2015;103(6):1384–96.

30. Kremer CT, Thomas MK, Litchman E. Temperature-and size-scaling of phytoplankton population growth rates: Reconciling the Eppley curve and the metabolic theory of ecology. Limnol Oceanogr. 2017;62(4):1658–70.

31. Listmann L, LeRoch M, Schlüter L, Thomas MK, Reusch TB. Swift thermal reaction norm evolution in a key marine phytoplankton species. Evol Appl. 2016;9(9):1156–64.

32. Masson-Delmotte V, Zhai P, Pirani A, Connors SL, Péan C, Berger S, et al. IPCC Climate change 2021: the Physical Science Basis. Contrib Work Group Sixth Assess Rep Intergov Panel Clim Change. 2021;2.

33. Zaiss J, Boyd PW, Doney SC, Havenhand JN, Levine NM. Impact of Lagrangian Sea Surface Temperature Variability on Southern Ocean Phytoplankton Community Growth Rates. Glob Biogeochem Cycles. 2021;35(8):e2020GB006880.

34. Ralston DK, Keafer BA, Brosnahan ML, Anderson DM. Temperature dependence of an estuarine harmful algal bloom: Resolving interannual variability in bloom dynamics using a degree-day approach. Limnol Ocean. 2014;59(4):1112–26.

35. Guillard RR. Culture of phytoplankton for feeding marine invertebrates. In: Culture of Marine Invertebrate Animals. Springer; 1975. p. 29–60.

36. Marañón E. Cell size as a key determinant of phytoplankton metabolism and community structure. Annu Rev Mar Sci. 2015;241–64.

