## Supplementary Information for "Multitrait diversification in marine diatoms in constant and warmed environments"

### Supporting Information

#### S1: Diatom cultures

Table S1: Details of the 6 *Thalassiosira* strains used in this study. Strain code, collection location and species ID from Bigelow culture collection, identification based on a BLAST search of the ITS 2 gene region sequences published in [1]

| Strain code | Collection location | Species | Identification based on ITS2 region |
|---|---|---|---|
| CCMP1010 | 37°N 65°W, Gulf Stream, between Bermuda and New York (approx.) | *T. weissflogii* | *T. weissflogii* |
| CCMP1050 | 32.966°N 117.251°W, Del Mar Slough, California USA | *T. weissflogii* | *T. weissflogii* |
| CCMP1587 | 6.08° S 106.79°E , Jakarta Harbor, Indonesia (approx.) | *T. weissflogii* | *T. weissflogii* |
| CCMP1059 | 19.665°N 156.034°W Aquaculture Pond, Oahu, Hawaii USA (approx.) | *Thalassiosira sp.* | *Cyclotella striata* |
| CCMP2929 | Unknown | *Thalassiosira sp.* | *T. weissflogii* |
| CCMP3367 | 44.9335°N 12.7005°E North Adriatic Sea | *T. pseudonana* | *T. pseudonana* |

#### S2: Trait measurements

Table S2: List of collected trait information for all experimental stages after seven days of preconditioning to 20°C and 24°C.

| Trait | Ancestors at start ($A_0$) | Reduced Selection (RS) | Ancestors at end ($A_{end}$) | Full Selection Control (FSC) | Full Selection High temp. (FSH) |
|---|---|---|---|---|---|
| Growth at 20°C | y | y | y | y | y |
| Growth at 24°C | - | - | y | y | y |
| Cell size at 20°C | y | y | y | y | y |
| Cell size at 24°C | - | - | y | y | y |
| Granularity at 20°C | y | y | y | y | y |
| Granularity at 24°C | - | - | y | y | y |
| Chl a at 20°C | y | y | y | y | y |
| Chl a at 24°C | - | - | y | y | y |
| Lipid content at 20°C | - | - | y | y | y |
| Lipid content at 24°C | - | - | y | y | y |
| ROS production at 20°C | - | - | y | y | y |
| ROS production at 24°C | - | - | y | y | y |
| Silicic acid uptake at 20°C | - | - | y | y | y |
| Silicic acid uptake at 24°C | - | - | y | y | y |
| POC content per cell at 20°C | y | y | y | y | y |
| PON content per cell at 20°C | y | y | y | y | y |
| POC/N ratio at 20°C | y | y | y | y | y |

#### S3: Growth rates during reduced selection

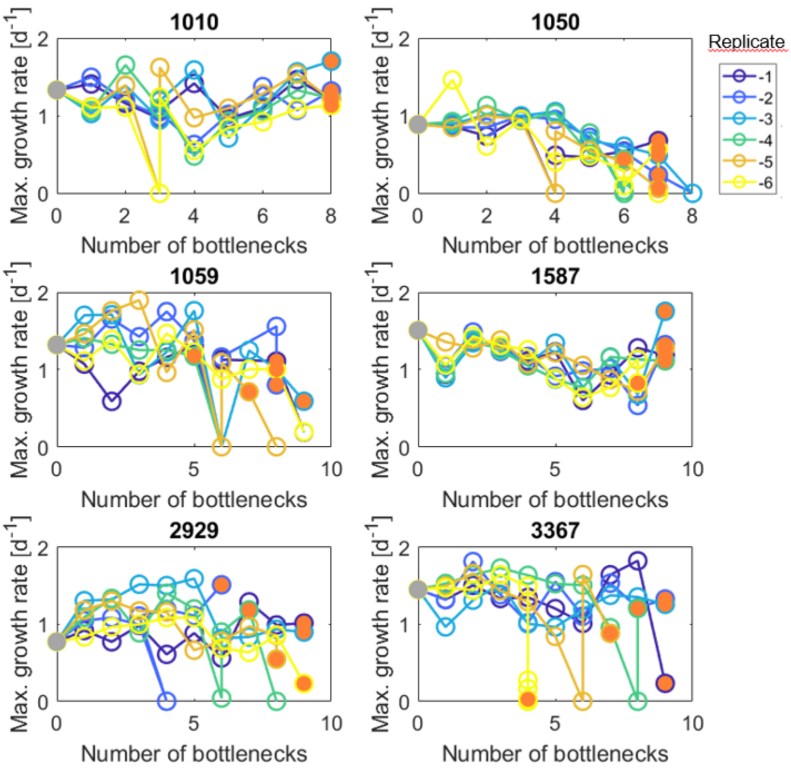


Figure S3: Maximum growth rates over the course of the reduced selection phase. Note that growth rates represent single replicates here. See Methods section for a detailed description of the bottlenecking procedure. Ancestral growth rate indicated by filled grey circles, final population used for the subsequent full selection phase indicated by orange filled circle. Vertical lines indicate that growth rates decreased to such a degree, that populations could not be maintained and the bottleneck had to be repeated with the samples from the previous transfer. Whereas strains 1050 and 1059 respond to the bottlenecking with a continuous decrease in maximum growth rates, strains 1010 and 1587 show only short-term fluctuations. Single replicates of 2929 and 3367 respond strongly to bottlenecking by reduced growth. By the end of the reduced selection, maximum growth rates decreased by 45%.

#### S4: Growth rates during full selection at control conditions

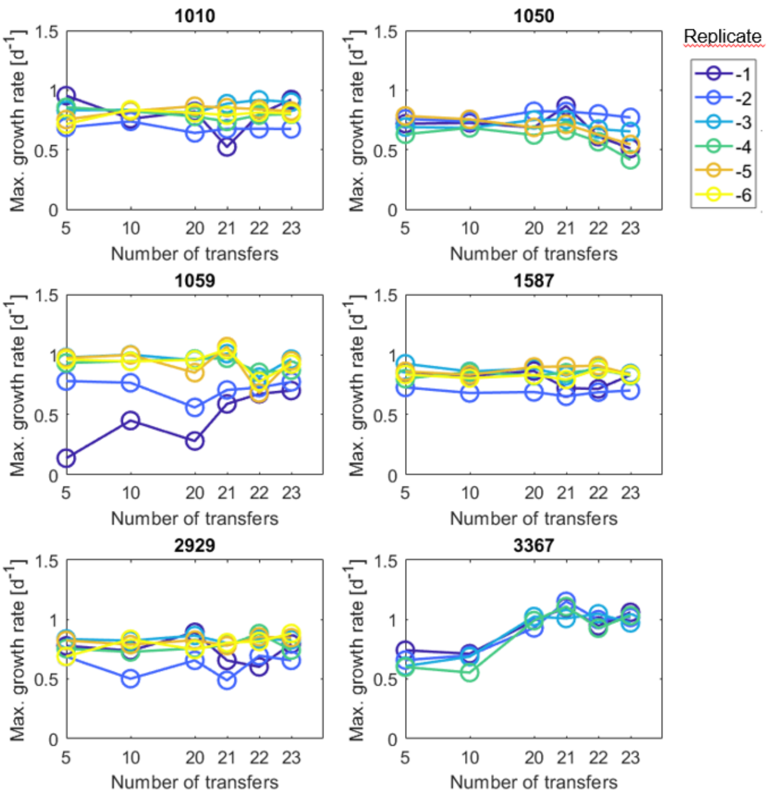


Figure S4: Maximum growth rates over the course of the full selection phase under control conditions. Note that growth rates represent single replicates here. Most strains showed stable growth rates over the course of the full selection phase. Replicates of strains 1059 and 3367, which experienced a strong reduction in growth during the reduced selection showed a recovery of high growth rates over the course of the full selection.

#### S5: Growth rates during full selection at high temperature conditions

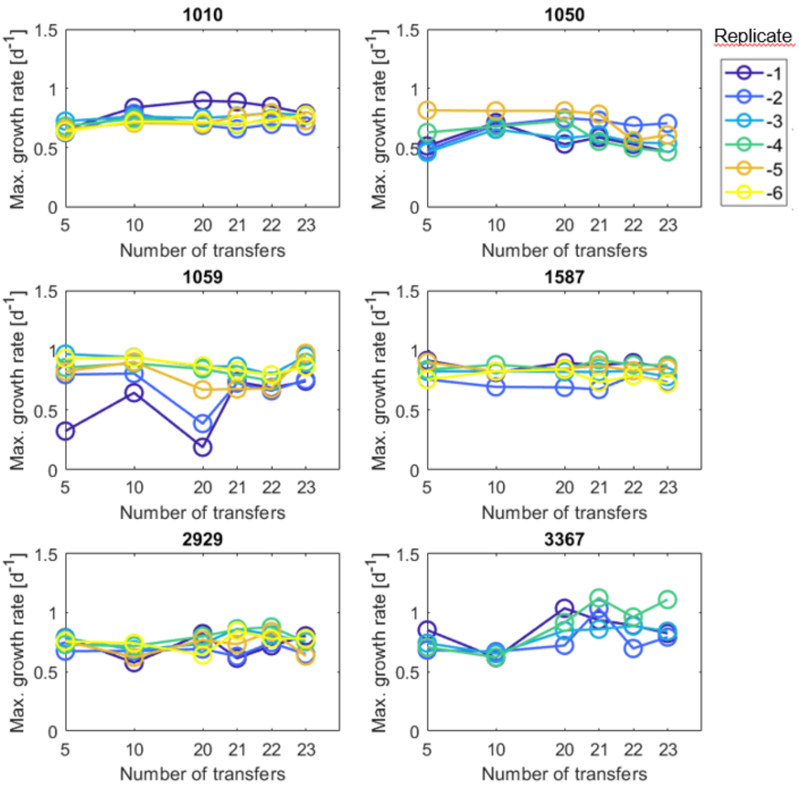


Figure S5: Maximum growth rates over the course of the full selection phase under high temperature conditions. Note that growth rates represent single replicates here. Growth rate patterns similar to full selection under control conditions.

**S6: Strain specific movements in trait-scape**

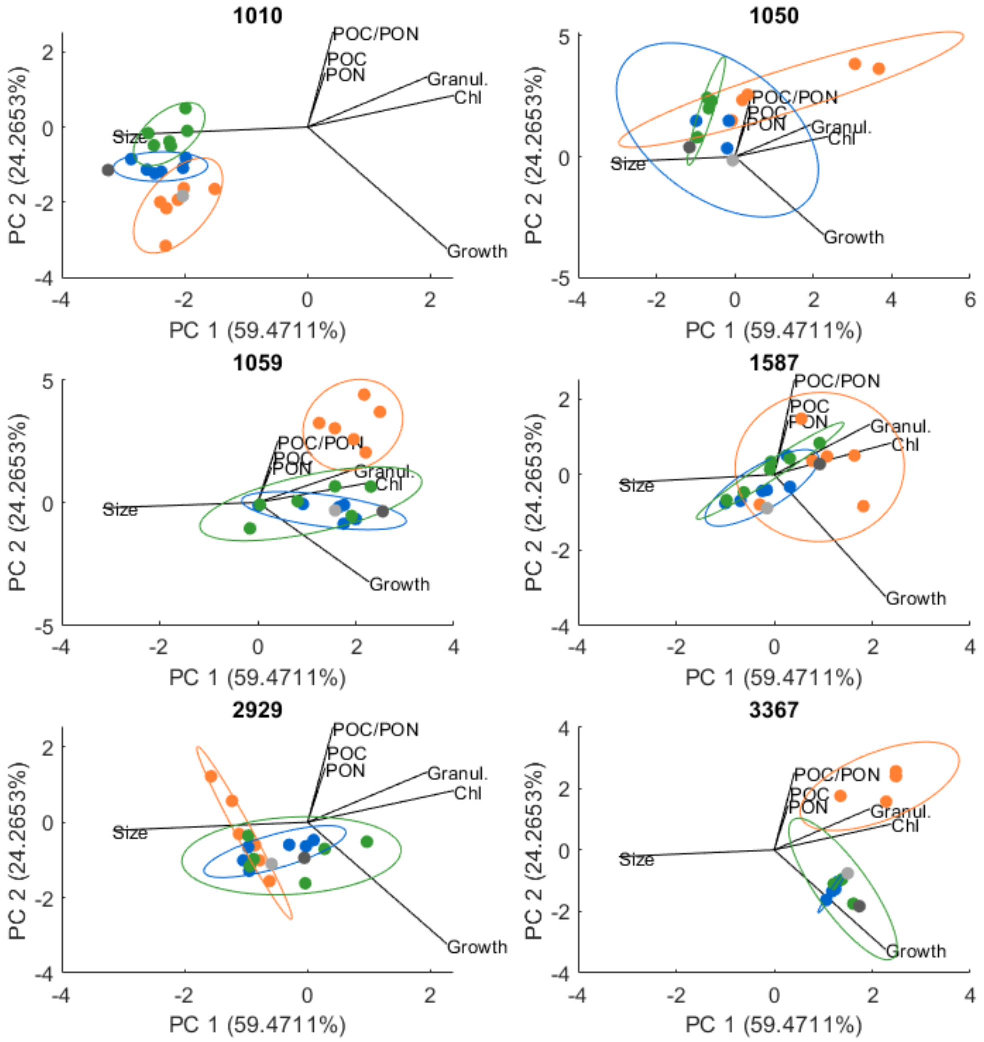


Figure S6: Movements in trait-scape of all strains from ancestors at the beginning of the experiment (light grey), to the location after reduced selection (orange), subsequent full selection under control (blue) and high temperature (green) conditions, as well as ancestors measured again at the end of the experiment (dark grey). Strains 1010, 1059 and 3367 show a clear divergence from ancestral trait values in response to reduced selection. The direction of the divergence in trait-scape is strain-specific. In response to full selection under control and under high temperature conditions, most populations recover locations in trait-scape that are close to their ancestors. Single populations cross into new areas in trait scape though, for example the high temperature evolved populations of 1010 moved to a new location in trait-scape.

#### S7: Single trait analyses

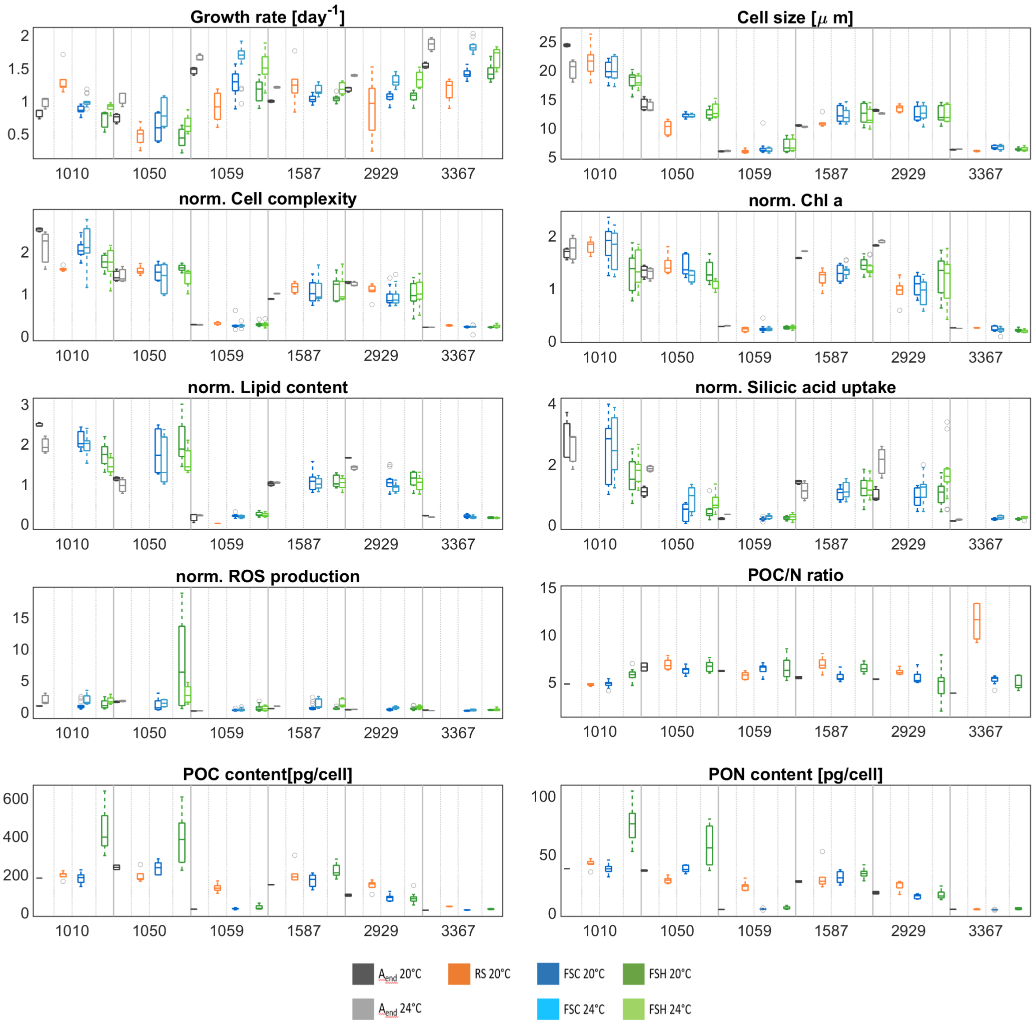


Figure S7: Trait boxplots across strains and experimental stages after acclimation to the 20°C control environment (dark grey, orange, dark blue, dark green) and the 24°C high temperature environment (light grey, light blue, light green). Experimental stages include ancestors at the end of the experiment (A, grey), reduced selection (RS, orange), full-selection under control conditions (FSC, blue) and full selection under high temperature conditions (FSH, green). Patterns in trait responses to reduced selection and full selection are stronlgy strain-specific.

#### S8: Pairwise trait correlations

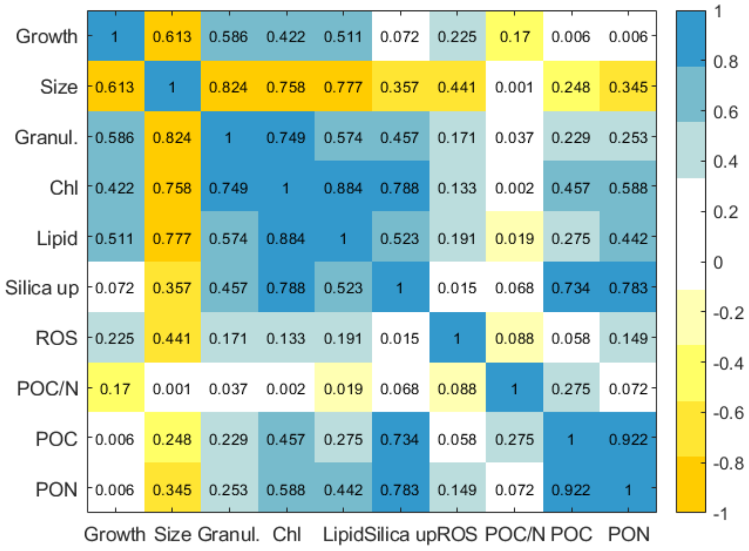


(a) Ancestral trait correlations

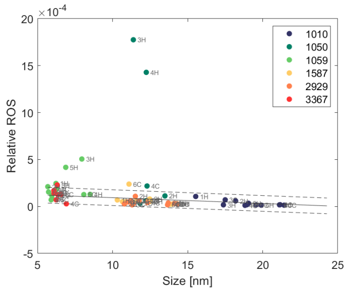


(b) Cell size vs. ROS

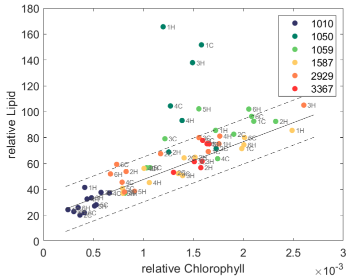


(c) Chlorophyll vs. lipid content

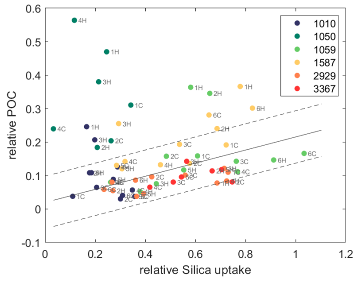


(d) Silica uptake vs. POC

Figure S8: (a) Ancestral trait correlations for all ancestors (Aend), including $r^2$ values. Most traits show a positive correlation, indicated by blue colours, only size is negatively correlation with all other traits. The POC/PON ratio shows little correlation with other traits, which is due to the strong correlation of POC and PON in all ancestral populations, leaving the POC/PON ratio at a value close to 1. (b)-(d) Example pairwise trait correlations. Shown are the ancestral trait correlations (solid line) and their 95% confidence interval (dashed lines), filed circles represent the measured trait correlations in FSC and FSH evolved populations. Whereas certain trait correlations are strongly conserved, such as cell size vs. reactive oxygen species, they can still allow for single strong outliers, where as other trait correlations such as silica uptake vs. POC content show overall more flexibility in trait correlations, but diversions from ancestral correlations remain modest.

#### S9: Outliers in trait-scape

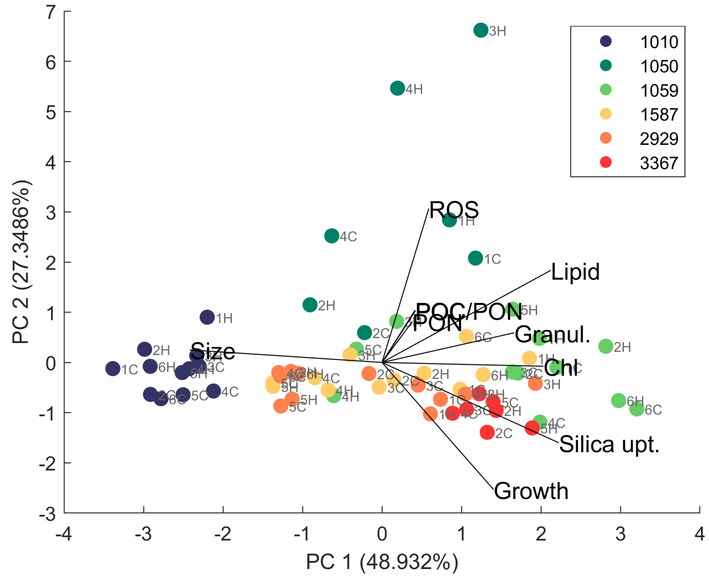


(a) Strains color-coded

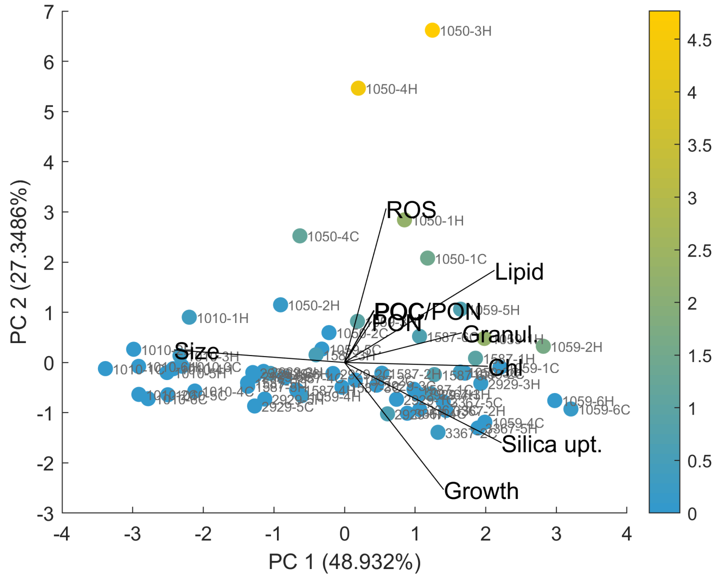


(b) Outlier score color-coded

Figure S9: The trait-scape including all traits of FSC and FSH population measured under control conditions. Outliers from ancestral trait correlations are indicated by colour. Strongest outliers are correlated with high values in ROS, POC and PON content, and lipid content.

#### S10: Outlier-growth analysis

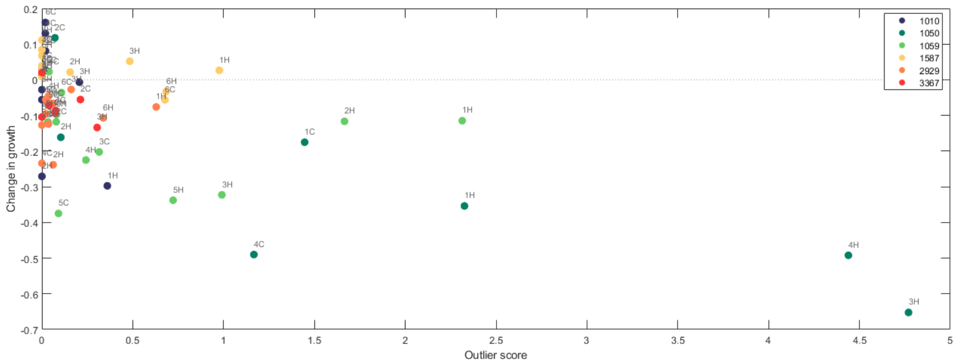


(a) Decreased growth of strong outliers

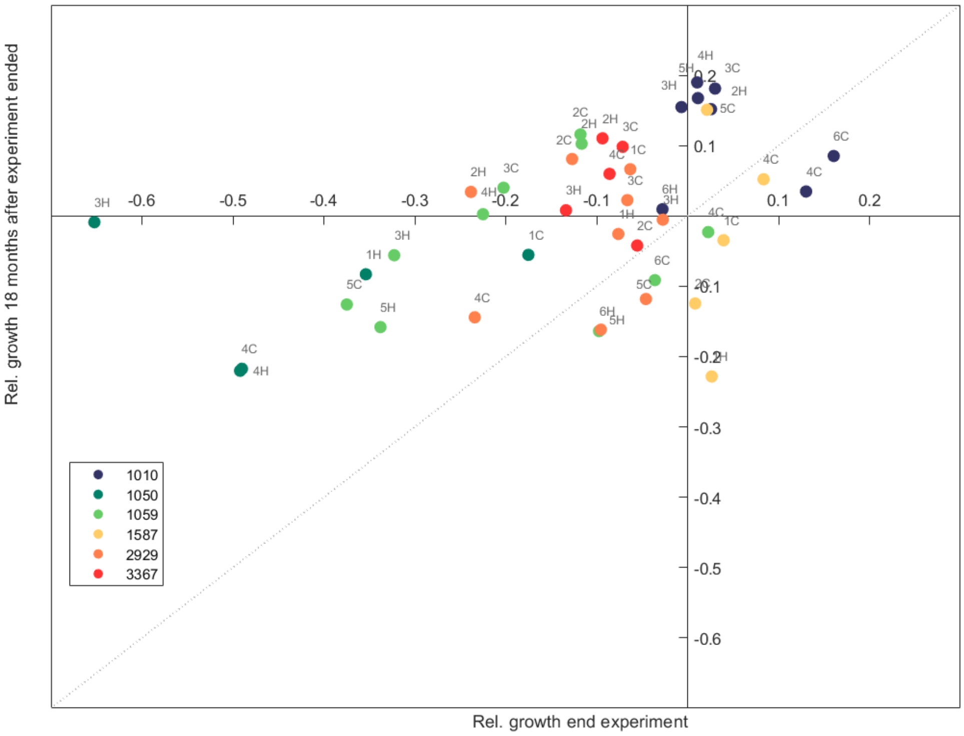


(b) Growth reanalysis

Figure S10: (a) Outliers vs. change in growth relative to ancestors. Strongest outliers show a decrease in growth rate in relation to their ancestors. (b) Relative growth rate at the end of the full selection phase (identical with y-axis in (a), and 18 months later. Most populations that showed a decreased growth rate after the full selection phase (e.g. 1050-3H and 1050-4H, recovered higher growth rates after the end of the experiment, indicating that their location in trait-scape did not represent a stable phenotypic trait combination.

#### S11: Trait boxplots of connected peaks

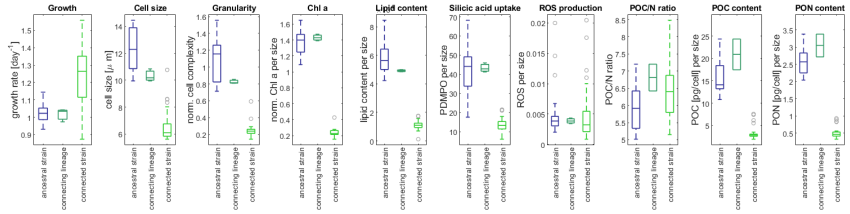


(a) 1587-1H

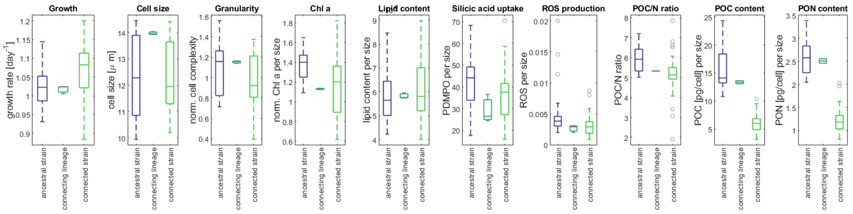


(b) 1587-5C

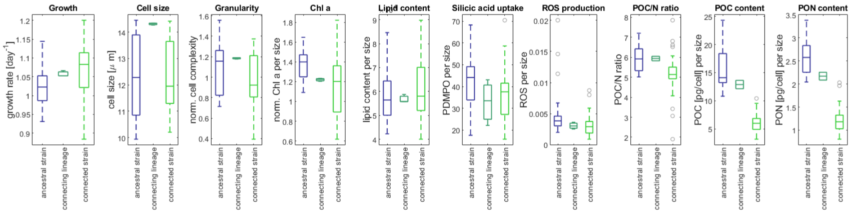


(c) 1587-5H

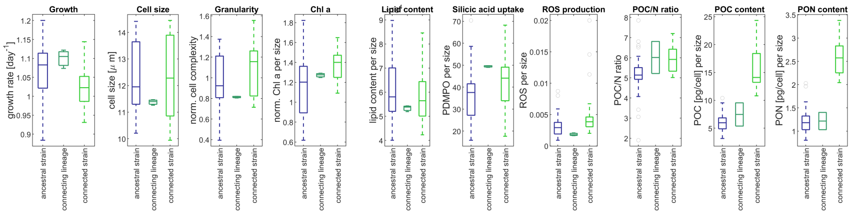


(d) 2929-1C

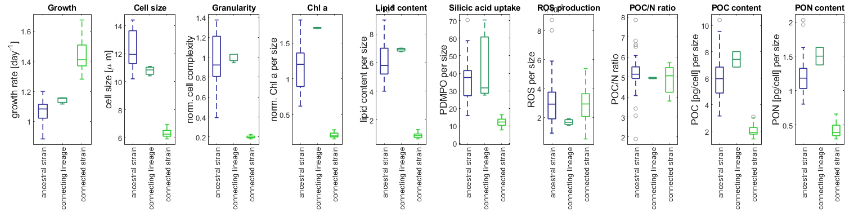


(e) 2929-3H

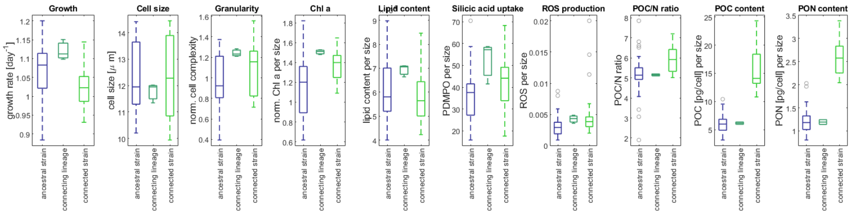


(f) 2929-4H

Figure S11: Boxplots of all connected peaks, illustrating the trait values of all evolved population from one ancestral strain (blue), the trait values of the lineage of that strain that moved into the trait-scape area of another, connected strain (dark green), and the trait values of all evolved populations of the connected strain (light green). See S12 for further analysis.

#### S12: Analysis of connected peaks

| Connecting lineage | Trait | Are ancestors different? | Movement outside ancestral range - past edge of box? | Movement towards connected strain? | Novel trait value? |
|---|---|---|---|---|---|
| **1587-5C** | growth | n | n | n | |
| | cell size | y | y | y | |
| | granularity | y | n | y | |
| | chl a | y | n | n | |
| | lipid content | y | y | y | |
| | silicic acid uptake | y | n | n | |
| | ROS | n | n | n | |
| | POC:PON | n | y | y | |
| | POC | y | n | n | |
| | PON | y | n | n | |
| 1587-1H | growth | n | n | n | |
| | cell size | n | y | n | * |
| | granularity | n | n | n | |
| | chl a | n | y | y | |
| | lipid content | n | n | n | |
| | silicic acid uptake | n | y | y | |
| | ROS | n | n | y | |
| | POC:PON | n | n | y | |
| | POC | y | n | n | |
| | PON | y | n | n | |
| 1587-5H | growth | n | y | y | |
| | cell size | n | y | n | * |
| | granularity | n | n | n | |
| | chl a | n | y | y | |
| | lipid content | n | n | n | |
| | silicic acid uptake | n | n | n | |
| | ROS | n | n | n | |
| | POC:PON | n | n | n | |
| | POC | y | n | y | |
| | PON | y | n | y | |
| 2929-1C | growth | n | n | n | |
| | cell size | n | n | n | |
| | granularity | n | y | n | |
| | chl a | n | n | n | |
| | lipid content | n | n | n | |
| | silicic acid uptake | n | y | y | * |
| | ROS | n | n | n | |
| | POC:PON | n | n | y | |
| | POC | y | n | n | |
| | PON | y | n | n | |
| **2929-3H** | growth | y | y | y | |
| | cell size | y | y | y | |
| | granularity | y | n | n | |
| | chl a | y | y | n | * |
| | lipid content | y | n | n | |
| | silicic acid uptake | y | n | n | |
| | ROS | n | y | n | * |
| | POC:PON | n | n | n | |
| | POC | y | y | n | * |
| | PON | y | y | n | * |
| 2929-4H | growth | n | n | n | |
| | cell size | n | n | n | |
| | granularity | n | y | y | * |
| | chl a | n | y | y | * |
| | lipid content | n | n | n | |
| | silicic acid uptake | n | y | y | * |
| | ROS | n | y | y | |
| | POC:PON | n | n | | |
| | POC | y | n | n | |
| | PON | y | n | n | |

Clearly staying with ancestor
Movement towards connected strain
Clearly moving towards connected strain
Novel trait value (unlike ancestor or connected strain)
Lineages in **bold:** majority of traits differ significantly

Figure S12: Characterization of connected peaks, based on trait values in S11. Trait values that are close to ancestor are marked in blue, trait values close to the connecting strain marked in green. Intermediate trait values between ancestor and connected strain are marked in grey. Trait values that neither resemble ancestral nor connected strain are marked in yellow. Even though some of the lineages developed trait values that resemble the connected strain in single traits, none of the lineages that were identified as connected peaks resembled the connected strain regarding the majority of traits. Instead, trait values that neither resemble the ancestor, nor the connected strain are common suggesting that the connecting lineages developed cryptic new phenotypes. The lineages of connected peaks thus do not necessarily match each other regarding trait values, but regarding a combination of trait values and trait correlations, that cannot be captured by analyzing single traits alone.

### References

[1] Phoebe A Argyle, Nathan G Walworth, Jana Hinners, Sinéad Collins, Naomi M Levine, and Martina A Doblin. Multivariate trait analysis reveals diatom plasticity constrained to a reduced set of biological axes. ISME Communications, 1(1):1–11, 2021.
